# Expression of HPV oncogenes from a single integration site in cervical cancer cell lines

**DOI:** 10.1101/2020.06.25.171959

**Authors:** Lulu Yu, Alexei Lobanov, Vladimir Majerciak, Sameer Mirza, Vimla Band, Haibin Liu, Maggie Cam, Zhi-Ming Zheng

**Affiliations:** Tumor Virus RNA Biology Section, HIV Dynamics and Replication Program, Center for Cancer Research, National Cancer Institute, National Institutes of Health, Frederick, MD 21702, USA; CCR Collaborative Bioinformatics Resource (CCBR), National Cancer Institute, Bethesda, MD 20892, USA; Department of Genetics, Cell Biology and Anatomy, University of Nebraska Medical Center, Omaha, NE 68198, USA

## Abstract

Integration of a virus genome into human chromosomal DNA is critical in viral carcinogenesis. In this report, we discovered that virus-host fusion transcripts are characteristically originated mainly from a single integration site in three cervical cancer cell lines, CaSki and SiHa cells with multiple HPV16 DNA integration sites and HeLa cells with multiple HPV18 DNA integration sites. The host genomic elements surrounding the integrated HPV genome are critical for efficient expression of the viral oncogenes. We found that HPV E6 and E7 are expressed by hijacking a host 3’ splice site and/or RNA polyadenylation signal for their production. The viral-host fusion transcripts may encode a chimeric viral-host fusion protein through alternative RNA splicing. One such E6* protein in SiHa cells was found to antagonize E6 function and knockdown of its expression further decreased p53 protein level and increased cell growth by promoting S phase entry. Together, our findings of the integrated virus genome expression only from a given integrated host genomic site will shed new light on possible application of precision medicine and the understanding of HPV carcinogenesis.

## Introduction

Human papillomavirus (HPV) is a non-enveloped, small DNA tumor virus. HPV replicates and assembles exclusively in the cell nucleus where the viral genome exists as an episomal chromosome. The HPV life cycle is highly linked to keratinocyte differentiation. Although HPV infections usually cause benign tumor, the high-risk HPVs, like HPV16 and HPV18, are causative biological agents of cervical cancers^1^, an increasing number of head and neck cancers and oropharyngeal cancers^2^. As the key event to carcinogenesis, HPV DNA integration into its host genome during its persistent infection is widely found in HPV-associated cancers^3–5^. HPV integration promotes carcinogenesis in various ways, the most important of which is to dysregulate the expression of viral oncogenes E6 and E7 that subsequently, impairs cell cycle control through inactivation of p53 and pRb, two cellular tumor suppressor proteins^6, 7^. The enhancement of E6 and E7 expression by HPV integration can be achieved often by disruption of the E1 or E2 region to alleviate E2 repression on E6 and E7 expression^8^. However, disruption of the E1 or E2 region also makes a viral early polyadenylation signal (PAS) inaccessible for polyadenylation of viral E6E7 bicistronic transcripts^9^ and thus, the E6E7 transcripts have to rely on an approachable host PAS in close proximity to the insertion site. It has been noted that viral E6E7 transcripts expressed from the integrated HPV DNA are more stable than the episomal viral transcripts, resulting in the increased E6 and E7 levels^10^. Despite over three decades of studies on HPV integration and gene expression, there is no report to date of how the integrated virus genome accomplishes its RNA splicing and polyadenylation to produce viral-host fusion transcripts at the integration sites. In the present study by mapping virus-host RNA splice junctions and identifying RNA polyadenylation cleavage sites of the RNA transcripts from the chromosomal HPV integration sites from three model cervical cancer cell lines, we discovered that only one integration site is most transcriptionally active regardless of the number of virus integration sites that exist in the host genome.

## Results and Discussion

### Identification of transcriptionally active integration sites by RNA-seq analysis of virus-host chimeric reads

To detect the transcriptionally active integration sites from HPV16-integrated CaSki and SiHa cells^11^ and HPV18-integrated HeLa cells^12^, we conducted extensive RNA-seq analyses and profiled the RNA expression pattern from each of three cell lines. Two types of paired-end reads were considered as virus-host “chimeric reads” in the current study: “chimeric paired reads” were the paired-end reads where one mate was fully aligned to the HPV16 or HPV18 genome (as defined by the aligner), while the other was fully aligned to the human genome; a “chimeric junction read or CJR” was the read that was partially aligned to the HPV16 or HPV18 genome and also partially aligned to the human genome from the junction with each overhang >=20 nts, while its mate was fully aligned to the virus or human genome (FIG 1A). Numbers of the chimeric reads mapped from CaSki, SiHa and HeLa cells are summarized in FIG 1B. We subsequently aligned these chimeric reads to the host genome to identify where the virus-host chimeric transcripts were transcribed in each cell line. As described below, we surprisingly discovered that the virus-host chimeric transcripts are expressed mainly from a single integration site in each examined cell line where multiple virus integration sites had been found^11, 13, 14^.

**FIG1.**
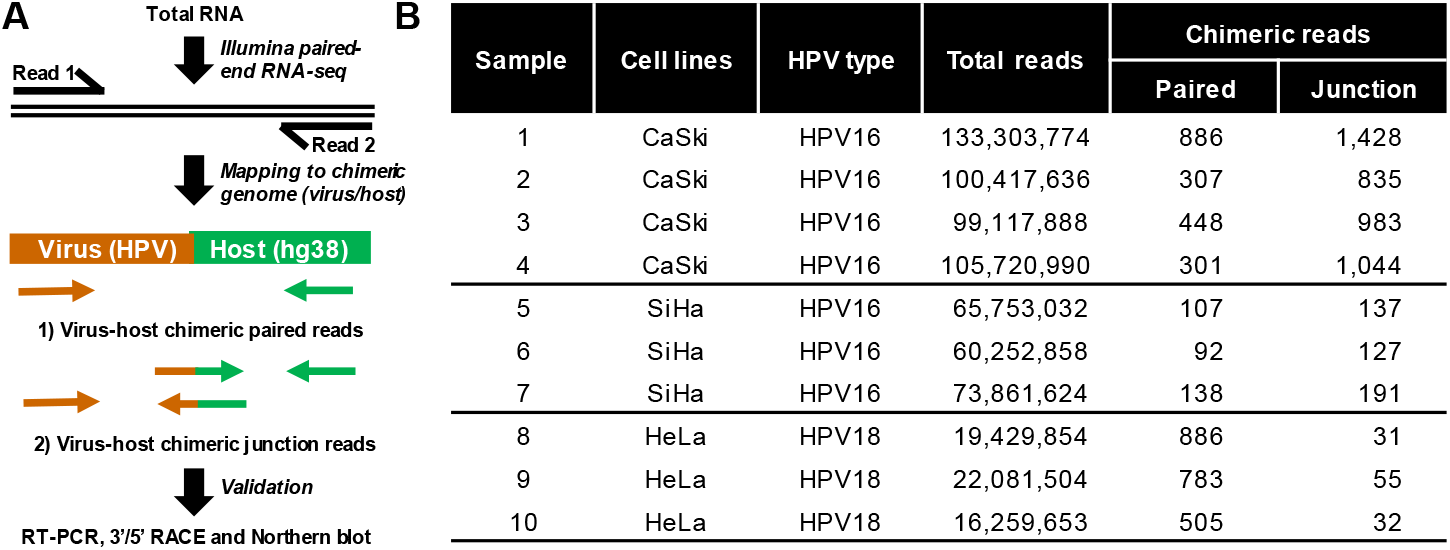
RNA-seq analysis of HPV integrations in CaSki, SiHa and HeLa cells. **A.** Diagram showing two principal types of chimeric RNA reads around a virus-host integration site. Paired-end reads from RNA-seq were mapped either to the HPV 16 or 18 genome and the human genome (hg38). The virus-host chimeric paired reads contain one read mate fully mapped to the virus and the other read mate fully mapped to the host. The virus-host chimeric junction reads (CJRs) are the reads mapped to the junction between virus and host RNA. RT-PCR, 3’ or 5’ RACE and Northern blot were used to verify the findings by RNA-seq. **B.** Number of chimeric reads from HPV16-positive CaSki and SiHa cells and HPV18-positive HeLa cells.

### Characterization of HPV16 insertion in CaSki cells reveals viral E6E7 expression primarily from a single integration site on chromosome 6

CaSki cells have been estimated to contain ~600 copies of the HPV16 genomic DNA per cell^11, 15^. To date, 44 different HPV16 integration sites have been identified in 11 different chromosomes (chr), including chr2, 3, 6, 7, 10, 11, 14, 19, 20, 21, and X, by whole genome sequencing (WGS)^13^. The number of identified integration sites may vary from different sequencing platforms with different depth coverages^14, 16^. Two integration sites in the chr6 intergenic region between CLIC5 and RUNX2 genes were confirmed from two recent reports^13, 14^, and could be further verified in this report by RNA-seq, PCR and Sanger sequencing (FIG 2). Analysis of this intergenic region showed that a repeat region from chr6 nt 45691285 to 45691466 appears unstable after HPV infection and tends to break^17^. We found by RNA-seq analysis that one integration site on the chr6 at nt 45691384 linking to HPV16 DNA at nt 3729 produces most of the virus-host CJRs (FIG 2A/B, also see FIG 2C diagram). As reported, the chr6 also breaks at nt 45691417 to link the linearized HPV 16 DNA at nt 2248 and the integration event leads to orientation reversion of the host genomic region^13, 14^. We confirmed both virus-host integration sites by PCR using a primer pair of B10 from the chr6 region and B4 from the HPV16 E1 region for the left integration site and a primer pair of F5 from the viral E2 region and F10 from the chr6 region for the right integration site (FIG 2D). By integration of the linearized virus DNA, the reversed chr6 region misses 33 nts from nt 45691417 to nt 45691384, presumably owing to digestion of the free DNA-chain ends by host exonucleases in the processing of virus DNA ligation to the host for the integration. Accordingly, we illustrated the integrated HPV16 DNA in the reversed chr6 region in CaSki cells (FIG 2C, top). In this diagram, HPV16 linearizes at two nick sites, one at nt 2248 and the other at nt 3729, each on the opposite strand of its double-stranded DNA to relax the supercoiled virus genome. DNA repair by filling each 5’ overhang end of the broken, relaxed linear virus genome leads to its integration into the chr6 at nt 45691417 for the E1 end and at the nt 45691384 for the E2 end in the orientation as indicated (FIG 2C). This integration allows viral early promoter P97 to transcribe viral early transcripts to utilize a host PAS AAUAAA at the chr6 nt 45691310, 74 nts downstream of the integration junction, for RNA polyadenylation and transcription termination, whereas the region upstream of the LCR lacks a visible promoter sequence and thus theoretically shouldn’t have transcriptional activity (FIG 2C, top). Subsequently, the RNA-seq reads were mapped to the chimeric host-virus genomic region and illustrated by IGV (FIG 2C, middle). The reads-coverage profile further confirmed the HPV16 integration and expression in the chr6 in the orientation we and others found^13, 14^ and exhibited the transcribed RNA splicing directions to use the host PAS for polyadenylation of the P97 transcripts (FIG 2C, middle), leading to production of three isoforms of the RNAs from alternative RNA splicing (FIG 2C, bottom). By 3’ RACE assays, we verified the usage of the host polyadenylation signal for RNA cleavage at nt 45691287 and polyadenylation of these transcripts from the integrated HPV16 genome in the CaSki cells (FIG 2E). The presence of three RNA isoforms in CaSki cells was verified by RT-PCR (data not shown) and by Northern blot analyses, with the isoform transcript-2 (E6*I) for E7 translation being the most abundant as reported^18, 19^. Together, these data indicate that the chr6 nt 45691384 site is the only integration site in CaSki cells for expressing abundant viral early transcripts from the integrated HPV16 DNA.

**FIG2.**
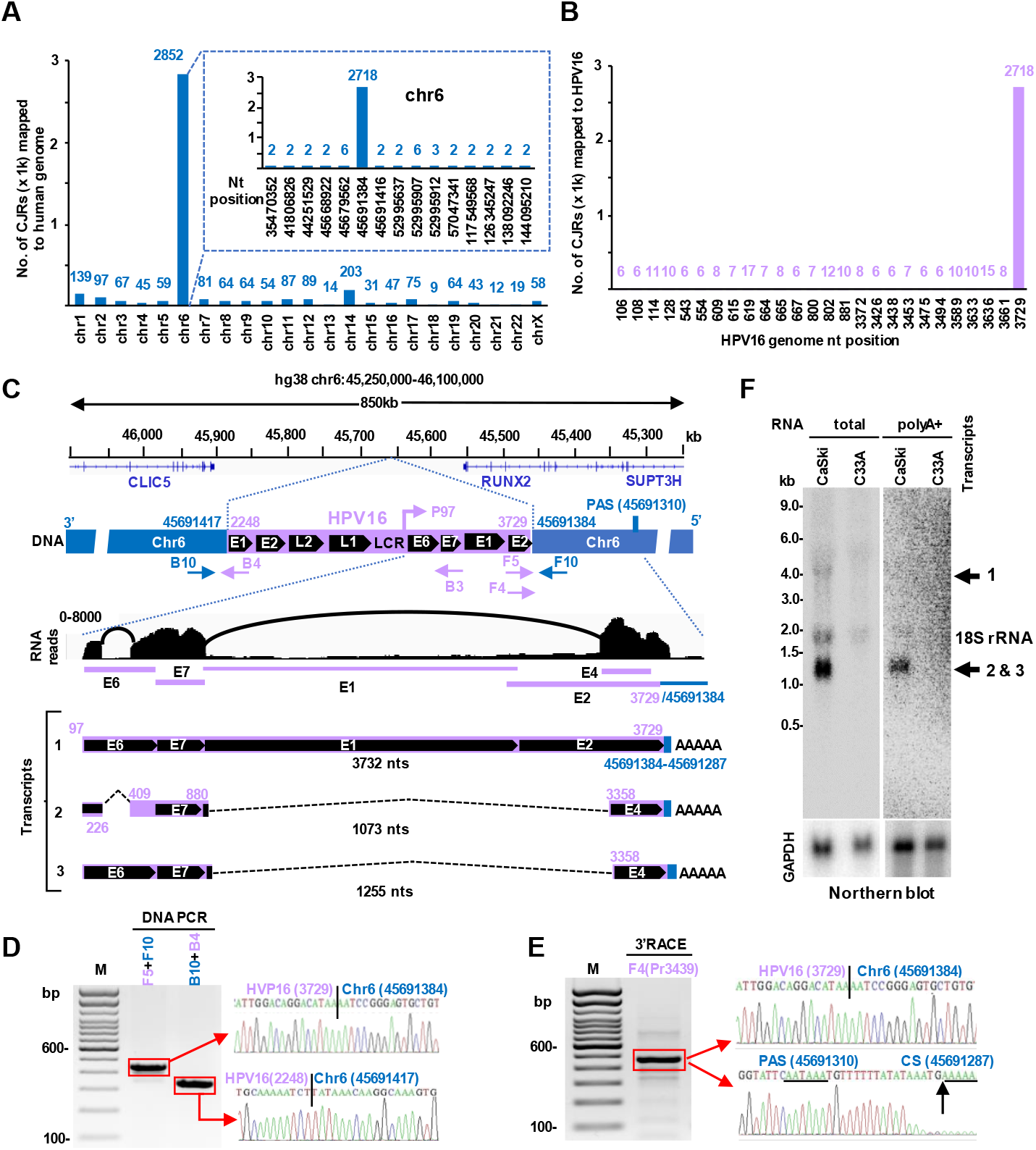
Transcription and RNA splicing from a single HPV16 integration site in CaSki cells. **A**. Distribution of the integration breakpoints in the human genome with the mapped CJRs>=6 in each chromosome, and CJRs>=2 in the chromosome 6 (chr6) at each integration site. **B**. Location of the integration breakpoints across the HPV16 genome with the CJRs>=6 at each detectable breakpoint. **C**. Diagram depicting HPV16 integration and expression in the chr6 in CaSki cells. Human chr6 breaks at nt 45691417 and nt 45691384 for HPV 16 DNA integration. By using the viral early promoter P97 and a host polyadenylation signal PAS at nt 45691310, the P97 primary transcripts undergo alternative RNA splicing to produce three RNA isoforms. Arrows below the chimeric genome are primers used for the validation. Total RNA-seq reads were mapped to the chimeric genome (HPV16:1-3729 /Chr6: 45691384-45691086) and visualized by IGV software. Three transcripts including pre-mRNA (transcript 1) and spliced mRNA transcripts 2 and 3 are shown. **D**. Validation of DNA breakpoints using the primer sets F5/F10 and B10/B4. The obtained PCR products (red rectangles) were gel purified and sequenced. Sequence readings are shown on the right. **E**. Mapping of virus-host chimeric transcript polyadenylation cleavage sites (CS) by 3’RACE using an HPV16 primer F4. The sequencing results containing the virus-host break point, PAS site, and cleavage site are shown on the right. **F**. Northern blot analysis of total CaSki RNA (10 μg) and poly(A)^+^ mRNA selected from 50 μg of total RNA using an antisense oligo probe B3 derived from HPV16 nt 855-836. Total RNA of HPV-negative C33A cells served as a negative control. GAPDH served as an RNA loading control. Major viral transcripts detected are shown on the right. 18S rRNA was from non-specific detection by this oligo probe.

### HPV16 E6E7 expression from chromosome 13 in SiHa cells by using a host 3’ splice site ~56 kb downstream of an integration site

SiHa cells have one to two copies of the integrated HPV16 genome per cell^11^. Unlike different HPV16 integration sites spreading over many chromosomes in CaSki cells, only two break sites, one at the chr13 nt 73214733 and the other at the chr13 nt 73513421, in SiHa cells were found by WGS in an intergenic, but reversely rearranged region between KLF5 and LINC00392 for HPV16 integration, where the linearized HPV16 genomic DNA in the E2 region at nt 3385 is linked to the nt 73214733 site and at nt 3133 to the nt 73513421^13, 14, 16^. This integration was most likely mediated by microhomology-mediated end joining (MMEJ)^20^ (Fig. S1). Our RNA-seq analysis showed the virus-host CJRs mapped to one of the two integration junctions from the HPV16 E2 region at nt 3133 to the chr13 nt 73513421, but majority of the CJRs were splice-junction reads derived from either a viral 5’ splice site at nt 226 in the E6 ORF or at nt 880 in the E1 ORF to a host 3’ splice site at the chr13 nt 73456962, further downstream of the nt 73513421 (Fig. 3A-C). We subsequently verified these splice junctions of virus-host RNAs by RT-PCR and sequencing of the RT-PCR products (Fig. 3D). By sequencing of PCR products amplified from total SiHa DNA, we confirmed, as reported^13, 14, 16^, the two integration sites with the inserted HPV16 DNA at nt 3385 to chr13 nt 73214733 in one site and nt 3133 to chr13 nt 73513421 in the other site (Fig. 3E). This integration takes place at the expense of losing 252 nts from the linearized virus genome breakpoint, probably by host exonuclease digestion of the free ends of linearized virus DNA.

**FIG3.**
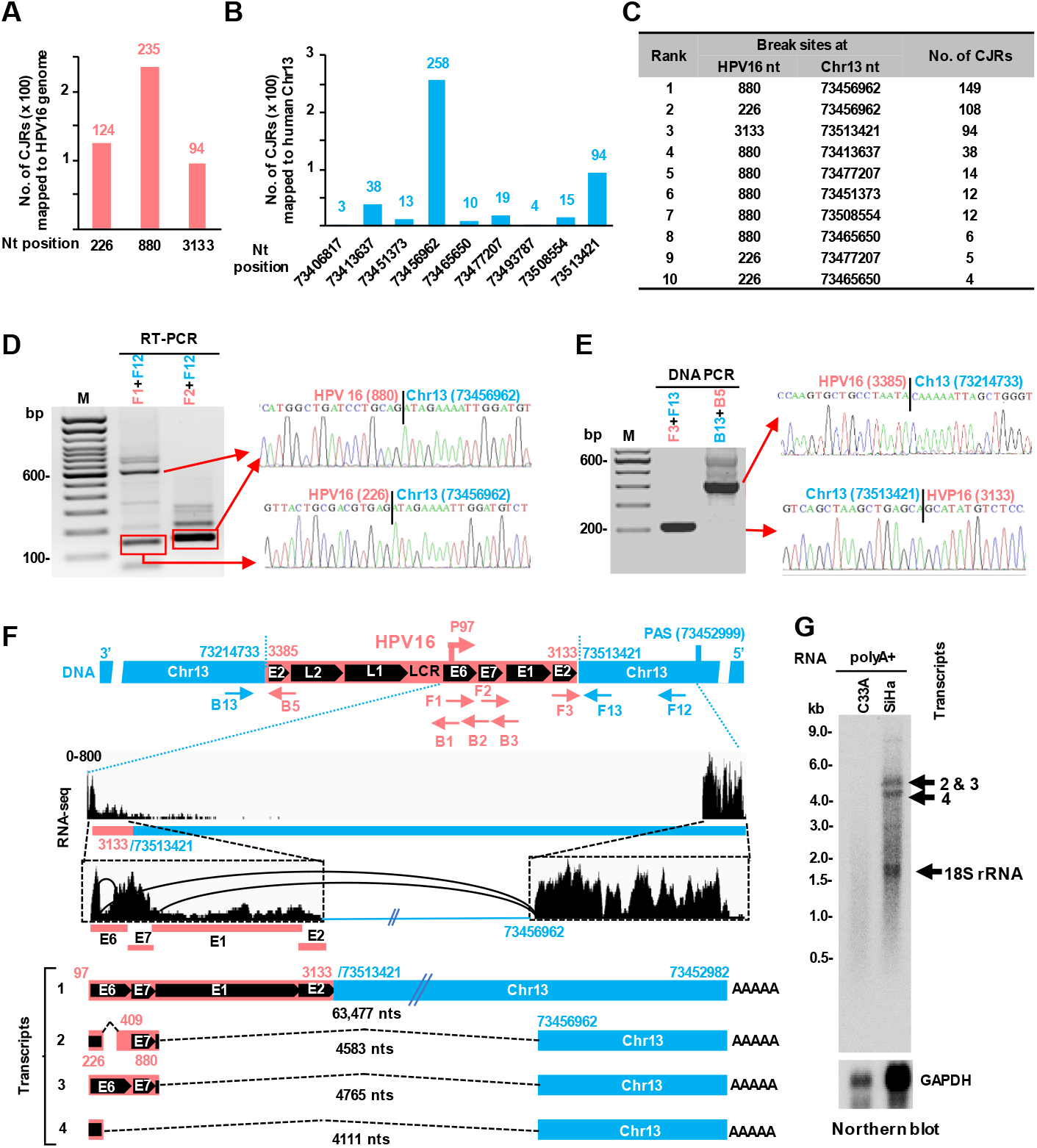
Identification of HPV16 RNA expression and splicing from a single integration site in HPV16-positive SiHa cells. **A** and **B**. Distribution of virus-host CJRs mapped to the HPV16 genome (A) and to chr13 (B) with the CJRs>=2. **C**. Distribution of top 10 CJRs mapped to the HPV16 and the chr13. **D**, Validation of two RNA splice junctions by RT-PCR using two primer sets F1+F12 and F2+F12 in panel F. Each RT-PCR product was gel-purified and sequenced with the sequence chromatographs showing the splice junction on the right. **E**, Two DNA integration break sites were verified by PCR using the primer set F3+F13 and B14+B5 in panel F and each gel-purified PCR product was sequenced with the sequence chromatographs showing the integration junction on the right. **F**. A diagram of HPV16 integration in the chr13, along with the reads-coverage and splicing of chimeric RNA isoforms from the integration site in SiHa cells. The illustration shows the HPV16 genome linearized at the E2 region from nt 3385 to 3133 and inserted to a reversed, rearranged locus of host chr13 between nt 73214733 and 73513421 positions. The primers used for the validation experiments are shown under the chimeric genome. Total RNA-seq reads were mapped to a chimeric genome (HPV16:3386-7906/1-3133/Chr13:73513421-73452495) and their distribution are visualized by IGV. The virus-host chimeric transcripts are transcribed only from a viral early promoter P97 of the integrated HPV16 genome and cleaved at chr13 nt 73452982 for polyadenylation. The chimeric transcripts are alternatively spliced to a host 3’ splice site at nt 73456962 to produce four isoforms of RNAs. **G**, Identification of the major transcripts in SiHa cells by Northern blot analysis using poly(A) selected mRNA from 100 μg of total RNA. A mixture of three probes (B1, B2 and B3) located in the HPV16 E6 and E7 regions were used for Northern blot analysis. RNA from HPV-negative C33A cells served as a negative control. Expression of GAPDH RNA served as a sample loading control.

Accordingly, we diagramed the integrated HPV16 genome in the orientation to the reversely rearranged chr13 region (Fig. 3F, top) and illustrated the transcription direction from the integrated HPV16 early promoter P97. This linearization and integration manner led the integrated HPV16 genome to lose its own PAS at nt 4215^9^ and inevitably select a host PAS for RNA polyadenylation of the P97 promoter-derived transcripts. Subsequently, we identified a canonical PAS AAUAAA at chr13 nt 73452999, ~60 kb downstream of the virus-host integration site at nt 73513421. By mapping of the total RNA-seq reads to the chimeric virus-host genomic region as visualized by IGV (FIG 3F, middle), the reads-coverage profile confirmed the HPV16 integration and expression in the chr13 and exhibited the transcribed RNA splicing directions to use the host PAS for polyadenylation of the P97 transcripts, including splicing of viral 5’ splice sites both at nt 226 and at nt 880 to a host 3’ splice site at nt 73456962, ~4 kb upstream of the host PAS (FIG 3F, middle). The RNA-seq reads-coverage profiles visualized by IGV also confirmed high reads-coverage from the host 3’ splice site to a polyadenylation cleavage site at nt 73452982, 17 nts downstream of the PAS at nt 73452999 (Fig. 3F). Because of the characteristics of this chimeric virus-host RNA structure and its alternative RNA splicing, four RNA isoforms were conceivably expected from viral P97 promoter-derived transcripts (FIG 3F, bottom) using the same PAS for RNA polyadenylation at the nt 73452982. By Northern blot analyses of poly-A^+^ SiHa cell RNA, we determined all three spliced isoforms (transcripts 2, 3 and 4) of the virus-host chimeric RNAs in the predicted size (Fig 3G), except the primary, unspliced precursor RNA transcript 1 (~64 kb). Among the three spliced transcripts, the transcript 2 is an E6*I RNA mainly for E7 translation^19, 21^, the transcript 3 with the E6 intron retention mainly encodes E6 protein and a possible E1^host fusion protein (FIG S2A), and the transcript 4 has the potential to encode an E6^host fusion protein of 55 aa residues from which 11 aa residues are encoded by host RNA portion (FIG. S2B).

### HPV18 expresses viral E6E7 in HeLa cells mainly from one integration site on chromosome 8

Each HeLa cell which contains 62-68 chromosomes has been estimated to have 20-50 copies of integrated HPV18 genome^22–24^. By using WGS, multiple HPV18 integration sites were recently identified at an intergenic, reversely rearranged region on chromosome 8 8q24.21^14, 16, 23, 24^. We re-analyzed the published data and illustrated the linearized, truncated HPV18 DNA integration in HeLa cells in each of three copies of chromosome 8, where a common integration site at nt 127218391 is linked to HPV18 nt 5736 in the L1 region at one end, but the other end varies among the three copies of chromosome 8, including nt 127229301 ligating to HPV18 nt 2497 in the E1 region, nt 127222007 to HPV18 nt 25 in the LCR region, or nt 127221122 to HPV18 nt 3100 in the E2 region^23, 24^. We realized that all integrated HPV18 DNA fragments are missing the viral early PAS and must use a host PAS for their expression. The illustrated integration site in each of the three copies of chromosome 8 could be further supported by WGS host/virus DNA CJR counts, with 421 CJRs for the common nt 127218391/nt 5736 junction, but 119, 195 and 33 CJRs, respectively, for individual junctions of nt 127229301/nt 2497, nt 127222007/nt 25 and nt 127221122/nt 3100 from each of the three copies of chromosome 8^14^. The latter two integration sites might express inefficiently because the integrated HPV18 nt 5736-nt 25 fragment does not have a viral promoter P55/102^25^ and the integrated viral nt 5736-3100 fragment gives low counts of the DNA CJRs. Analysis of the surrounding DNA sequences in the functional HPV18 integration sites (nt 5736 to nt 2497) in HeLa cells showed the integration event most likely mediated by MMEJ^20^ (Fig. S1).

By mapping our RNA-seq raw reads to HPV18 and hg38, we identified the top four most common virus/host CJRs derived from nt 2497/nt 127229301, 929/127229130, 24/127218810 and 5736/127218391 (FIG. 4A-4C). Two, nt 2497/nt 127229301 and nt 5736/nt 127218391, represented the integration junction RNA reads in consistence with the mapped integration sites in the Chr8 detected by WGS^23, 24^, and the other two represented the splice junction reads of virus-host chimeric RNAs. By using total DNA extracted from HeLa cells, we further verified the two common integration junction sites by PCR and sequencing of the PCR products (Fig. 4D). Accordingly, we depicted the integrated HPV18 genome in the orientation to the reversely rearranged chr8 region and the transcription direction from the integrated HPV18 early promoter P55/102^25^ (Fig. 4E, top). The mapped RNA-seq reads distribution around the chr8 integration site was visualized by IGV and showed RNA splicing direction, activation of a weak promoter (P127219268) within the reversed intron 1 of host lncRNA CCAT1^26^ upstream of the integrated HPV18 DNA, and utilization of an expected host PAS downstream of the integrated viral DNA for polyadenylation of the chimeric virus-host RNA transcripts (Fig. 4E, middle). We verified the RNA cleavage site for RNA polyadenylation at chr8 nt 127228559, 17 nts downstream of the PAS at nt 127228576 and all RNA splice junctions by 3’ RACE and sequencing (FIG 4F-4G). Because of alternative RNA splicing, five isoforms of the RNA transcripts including a unspliced pre-mRNA transcript 1 are predicted and three of them are E6* I mRNA fusions to encode viral E7^19, 27^, except the transcript 5 being a minor isoform encoding viral E6 (Fig. 4E, bottom). Northern blot analysis of HeLa cell total or poly-A^+^ RNA by using an antisense viral probe confirmed the major transcripts of the expected size (FIG 4H).

**FIG 4.**
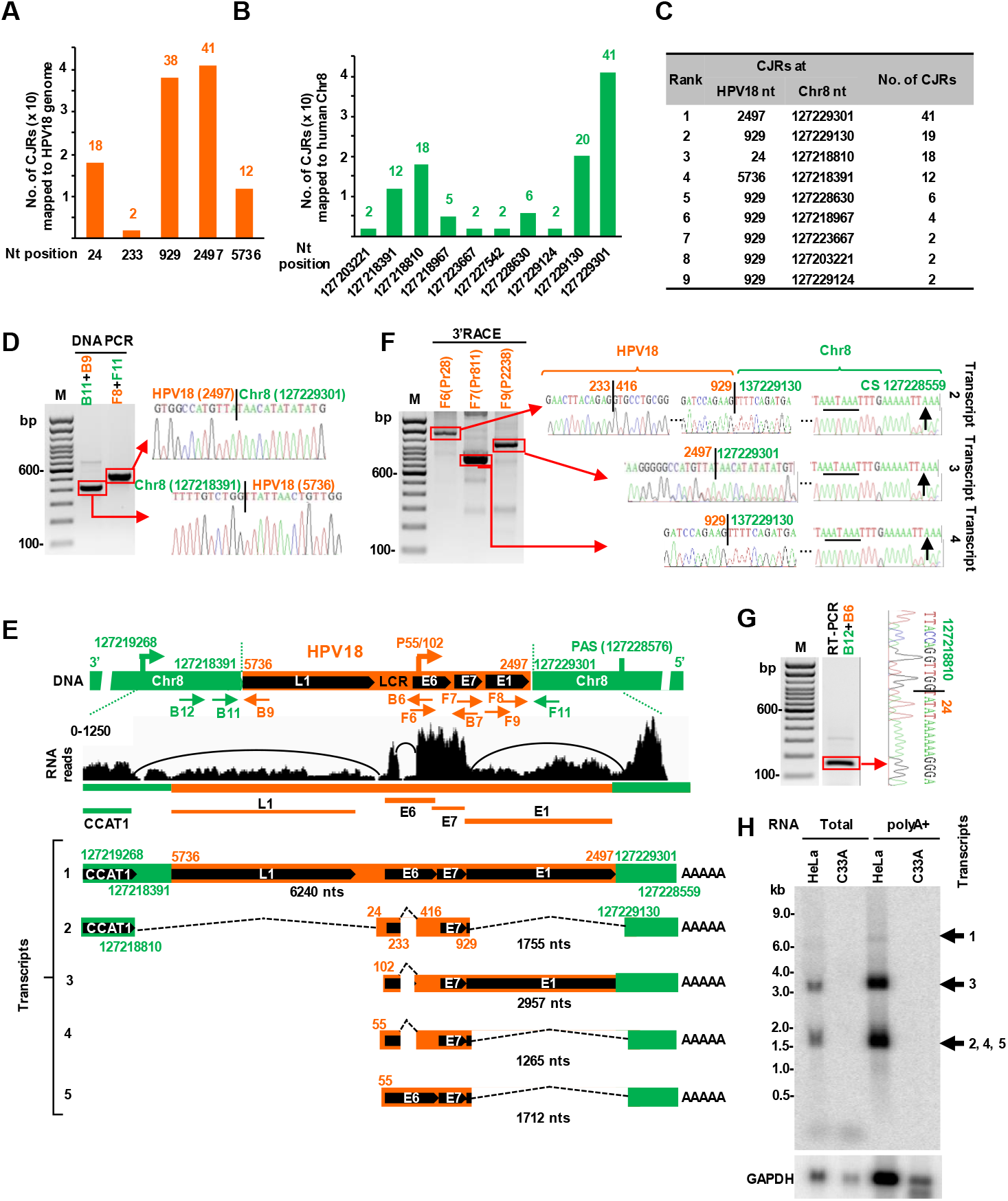
HPV18 transcription and RNA splicing from a single integration site in HPV18-positive HeLa cells. **A & B.** Mapping of virus-host CJRs to HPV18 genome (A) and human chr8 (B) with CJRs>=2. **C.** Distribution of top 9 virus-host CJRs identified by RNA-seq. **D.** Verification of two integration junction sites in HeLa cells by PCR using primer set B11+B9 and F8+F11 in panel E. **E.** Illustration of HPV18 integration and expression from a chr8 integration site in HeLa cells. A linearized, truncated HPV18 genome in size of 4618 nts is inserted to a reversed, rearranged chr8 region by ligation of the viral DNA end at nt 5736 in the L1 region to chr8 nt 127218391 and the other viral DNA end at nt 2497 in the E1 region to chr8 nt 127229301at the expenses of losing a ~3.2-kb viral DNA fragment and a ~10.9-kb host DNA fragment. As one of the two integrated DNA junctions happens in the reversed intron 1 region of an antisense transcript lncRNA CCAT1, this integration disrupts the CCAT1 expression. However, the CCAT1 promoter becomes available, in addition to the HPV18 early promoter P55/102, for the expression of integrated HPV18. Arrows below the chimeric genome are the primers used for various validations. RNA-seq reads-coverage along the HPV18 integrated chr8 region was visualized by IGV, illustrating RNA transcription, RNA splicing direction and polyadenylation to produce five isoforms of RNA transcripts, with the RNA transcript 1 as a unspliced pre-mRNA. **F.** Detection of HPV18 polyadenylation cleavage sites (CS) by 3’ RACE using an HPV18 specific-primer F6, F7 or F9. The splicing or integration junction and PA cleavage site (black arrow) identified by Sanger sequencing are shown on the right. **G.** RT-PCR validation of a viral-host fusion RNA splice junction using a primer set B12+B6 with the sequencing result on the right. **H.** Northern blot detection of the major RNA isoforms transcribed from the identified integration site in HeLa cells. Total RNA at 10 μg and poly(A)-selected mRNA from 100 μg of total RNA were examined by Northern blotting using an antisense HPV18 probe (B7). The corresponding RNA preps from an HPV-negative cervical cancer cell line C33A served as negative controls, with GAPDH RNA serving as a sample loading control.

### Expression of viral E6 and/or E7 with point mutations and functional identification of viral-host fusions from the integration sites

By analyzing the distribution of mapped RNA-seq reads in the integrated virus genome in all three cell lines by IGV, we identified various point mutations when aligned with the corresponding reference HPV genomes (Fig. S3A-C). Data are consistent with the reported observations^28^, except the nt 2609T in CaSki cells. These mutations in the E6 and E7 ORF regions could be silent or missense mutations. The missense mutations lead to a subsequent change in the amino acid residues of E6 and/or E7 proteins (Fig. S3D-F), of which their functional consequences remain to be studied.

Given that a host PAS downstream of the virus integration site in a host chromosome region is essential for efficient expression of the integrated HPV genome and often reachable by alternative RNA splicing into the host region, we sought out for possible viral-host fusion proteins expressed from the virus integration sites and predicted four putative viral-host fusion proteins in size of >50 aa residues from four different fusion RNA transcripts identified in SiHa and HeLa cells (Fig. S2), but none from CaSki cells. The fusion transcript 1 transcribed from a host promoter in HeLa cells exhibited the RNA reads-coverage in the truncated L1 region, but this integration creates a frameshift in the truncated L1 ORF and thus does not encode L1 protein. The L1 PAS signal in this transcript was also inactive in our 3’ RACE analyses (Data not shown). Notably, a large truncated E1 protein in size of 528 aa residues could be predicted from the transcript 3 (Fig. 4E) in HeLa cells by using a stop codon right from the host junction (Fig. S2D). However, the transcript 3 in HeLa cells is an E6*I mRNA which functions mainly as an E7 mRNA^19, 27^ and unlikely translates efficiently the truncated E1 because the E1 start codon AUG is only 7 nts apart from the E7 stop codon upstream, which limits a translating ribosome from translational re-initiation^19, 29, 30^. We subsequently investigated the potential function of the fusion protein 2 encoded by the fusion transcript 4 derived from the integrated HPV16 E6 region in SiHa cells (Fig. S2B). The N-terminal 41 aa residues of this fusion protein in size of 55 aa residues are derived from the N-terminal half of E6 protein and shorter only by two aa residues from HPV16 E6*I protein^21^. By using a siRNA specifically targeting the splice junction of transcript 4 (Fig. 5A), we efficiently knocked down the expression of transcript 4 in SiHa cells (Fig. 5B-C). The specific KD of transcript 4 led to decrease the level of p53 by Western blot analysis (Fig. 5D) and increase the cell growth in a time-dependent manner (Fig. 5E). This was expected since viral E6*I modulates the E6-directed degradation of p53 and suppresses transformed cell growth^31, 32^. Consistent with these observations, cell cycle analysis by flow cytometry showed that siRNA-mediated specific reduction of transcript 4 significantly promotes cell entry into S phase.

**FIG 5.**
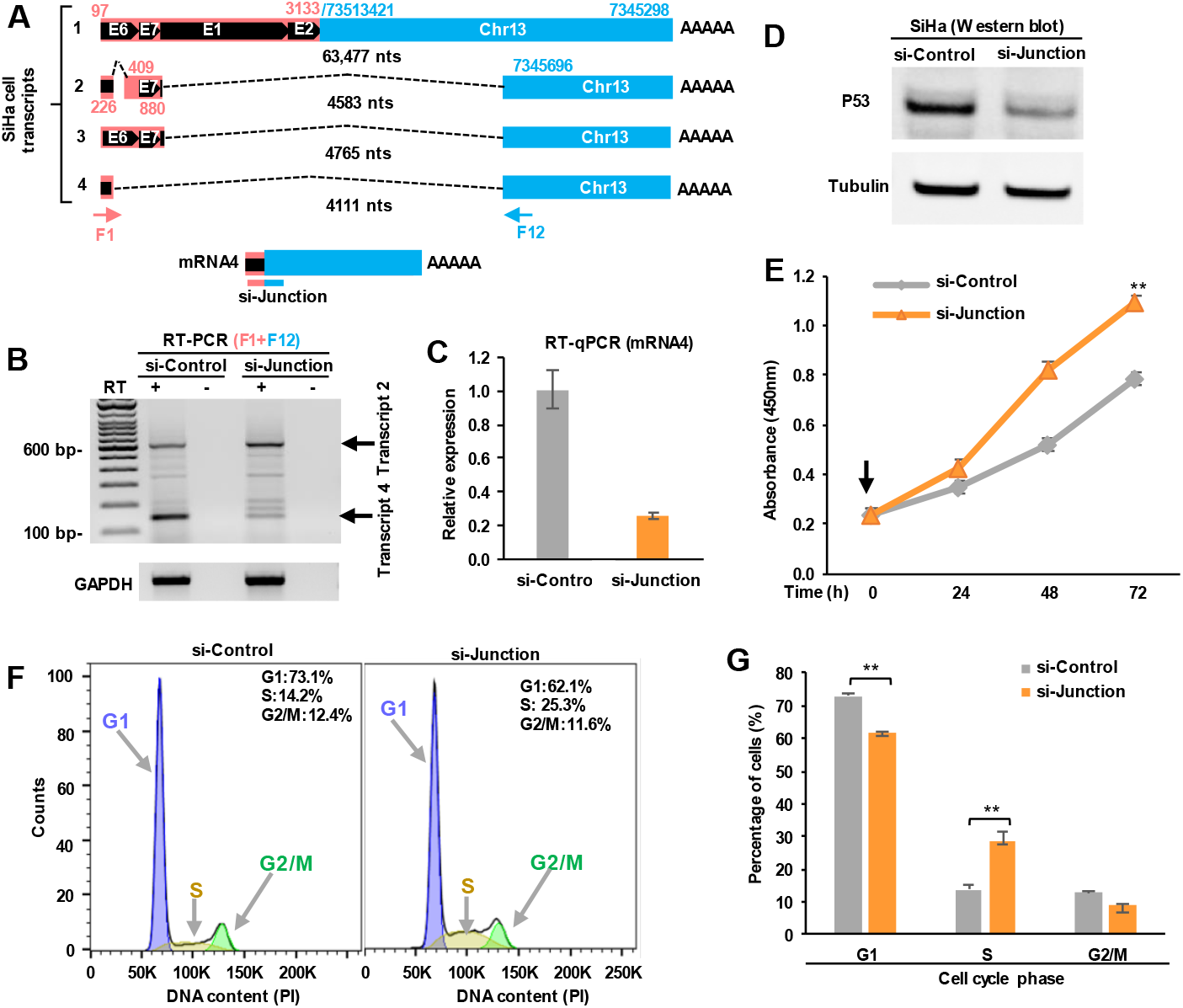
Reduction of a viral-host fusion transcript 4 in SiHa cells promotes cell proliferation. **A**. Major viral-host fusion transcripts in SiHa cells. Arrows are the primers used for RT-PCR detection of the fusion transcripts. A synthetic siRNA specifically targeting the transcript 4 (si-Junction) was custom-designed. **B**-**D**. Reduction of the transcript 4 expression in SiHa cells led to reduction of p53 protein. SiHa cells were harvested for total cell protein and RNA preparation 48 h after siRNA transfection. Knockdown efficiency of the transcript 4 was evaluated by RT-PCR using a primer set of F1+F12 (B). Expression of the transcript 2 and GAPDH served as a sample loading control. Knockdown efficiency of the transcript 4 was also evaluated by RT-qPCR using a virus-host specific junction TaqMan probe (C). Effect of transcript 4 knockdown on p53 expression was examined by Western blot with □-tubulin serving as a sample loading control (D). **E.** Enhancement of SiHa cell proliferation by knockdown of transcript 4 expression. SiHa cells at 24 h after plating were transfected with 40 nM of siRNA (arrow, si-Junction or si-Control) and the cell proliferation was examined at the indicated time (h) after siRNA transfection by averaged cell number mean ± SE from three experimental repeats. **, p<0.05 by unpaired, two-tailed Student’s *t* test. **F & G.** Reduction of the transcript 4 promotes cell entry into S phase. Flow cytometry analysis for cell cycle distribution after siRNA (si-Junction or si-Control) knockdown. SiHa cells at 24 h after plating were transfected with 40 nM of siRNA and harvested for flow cytometry analysis 48 h after siRNA transfection. One representative of three separate experiments is shown (F). Percentage of cells in the indicated phases of the cell cycle after siRNA knockdown of transcript 4 are shown in bar graphs (G). Data are mean ± SD; n=3. **, p<0.05 by unpaired, two-tailed Student’s *t* test.

### Conclusion

HPV integration leading to increased expression of E6 and E7 oncogenes is essential for HPV carcinogenesis. Early studies were limited to the detection of the integrated HPV DNA, roughly map the HPV integration sites, or estimate the copy numbers in host chromosomes in cervical cancer cell lines and tissue samples^33–36^. The restraints to precise mapping of the genome-wide HPV integration sites were alleviated recently by whole genome sequencing^13, 14, 23^. To date however, many questions remain open. What dictates the expression activity of the integrated HPV DNA? Are viral E6 and E7 oncogenes expressed from every integration site on different host chromosomes? Are viral-host fusion proteins expressed from the integrated HPV DNA? In this study, we demonstrated that only one HPV integration site out of multiple HPV integration sites is transcriptionally active in all three examined cervical cancer cell lines. Our findings are consistent with a reported study that only a single active transcription site was detectable by fluorescence in situ hybridization^37^ in CaSki cells containing more than 40 HPV16 integration sites on 11 different chromosomes^13^. We found that the efficient expression of viral oncogenes and viral-host fusion transcripts depends on the integrated HPV DNA in a given host genome region where other essential elements are available for RNA processing of the viral transcripts.

Unlike retrovirus site-specific integration which is catalyzed by a viral integrase and is an obligatory replication step^38–40^, the integration of HPV DNA into host chromosomes is a dead-end of the viral life cycle and a nonspecific, randomized event although the preference to host DNA fragile sites, intergenic regions and transcriptional active sites have been proposed^41–43^. Our comprehensive findings of the viral-host fusion transcripts and their transcription and RNA processing patterns in CaSki, SiHa and HeLa cells indicate that HPV integration in all three cell lines, albeit on different chromosomes, all of which led to the host genome instability at the integration site where the inserted HPV DNA is linked by a re-arranged, reversed host DNA on each side. We acknowledge that the characteristic features of HPV integration sites must be in favor of viral E6 and E7 expression and production of new viral-host fusion transcripts that may or may not encode a fusion protein, therefore resulting in different oncogenic potentials in different infected cells for clonal expansion. From our observations from multiple cell lines, it appears that a byproduct of HPV integration depends on the different linearized forms of HPV DNA, which can efficiently utilize both a viral promoter as well as a host polyadenylation signal at the integration site of the host genome to produce a fusion transcript. Altogether, our results shed additional novel mechanisms on how HPV integration and production of viral-host fusion transcripts can contribute to the development of HPV-associated cancers.

## Material and Methods

### Cell lines and siRNAs

HPV16-positive cervical cancer cell lines, CaSki and SiHa cells, and HPV18-positive cervical cancer cell line, HeLa cells were obtained from ATCC (Manassas, VA). HPV-negative cervical cancer cell line, C33A cells (mutant p53, mutant pRB) from ATCC, was used as an HPV-negative cell control. All cell lines were maintained in Dulbecco’s modified Eagle’s medium (DMEM) (Thermo Fisher Scientific, Waltham, MA) with 10% fetal bovine serum (FBS, GE Healthcare, Logan, UT) at 37°C under a 5% CO_2_ atmosphere.

To target the viral-host fusion RNA transcripts, a custom-designed synthetic siRNA specifically targeting the virus-host junction of the fusion transcript 4 in SiHa cells was (Table S1) was purchased from Dharmacon (Lafayette, CO). Non-targeting control siRNA (Dharmacon, # D-001210-01) served as a negative control. SiHa cells at 24 h of cell passage were transfected with 40 nM of siRNA by LipoJet In Vitro Transfection Kit (Ver. II) (SignaGen Laboratories). Total protein extracts and total RNA were prepared 48 h after the transfection.

### Total RNA sequencing and bioinformatics analysis

The total RNA extracted by TriPure reagent (Roche, Germany) was used for construction of RNA-seq libraries followed by Illumina sequencing. Sequencing libraries were constructed following Illumina Stranded Total RNA protocol (Illumina, RS-122-2201). Paired-end 150-bp read length sequencing with a depth of 100 million reads per sample was performed on the HiSeq 2500 sequencer according to the manufacturer’s instructions (Illumina) for CaSki cells. For SiHa cells, the length of the paired-end read is 75-bp with a depth of 75 million reads per sample. For HeLa cells, the length of the paired-end read is 50-bp with a depth of 25 million reads per sample. Adapters and low quality bases were trimmed by CutAdapt v1.18 with the following parameters: cutadapt --pair-filter=any --nextseq-trim=2 --trim-n -n 5 -O 5 -q 10.10 -m 35:35. Reference sequence of HPV16R used in this study was identical to HPV16 REF.1 (gi|333031) (https://pave.niaid.nih.gov), except G substitution of A at nt 2926 position. HPV18REF.1 (gi|60975) (https://pave.niaid.nih.gov) and human genome (GRCh38, hg38) were also used to create chimeric HPV+hg38 genomes for reads mapping. Reads were aligned using STAR v 2.5.2b aligner with default setting^44^. The paired reads fully aligned to the human or HPV16/18 genomes were removed, and the remaining “chimeric read pairs” were used in the follow-up analysis. We identified two types of “chimeric reads”. Chimeric paired reads were defined as that one mate was fully aligned to HPV genome and the other mate was fully aligned to human genome. Chimeric junction reads were the junction reads partially aligned to both HPV and human genomes with a threshold of minimal overhang >=20 nts from the junction to each genome, with a corresponding mate fully aligned to HPV or human genome. STAR-generated “Chimeric.out.junction” files were used to calculate the number of chimeric reads. The uniquely mapped reads were then uploaded into the Integrative Genomic Viewer (IGV) program (https://software.broadinstitute.org/software/igv/) to visualize reads-coverage profile along with individual genomes. A Sashimi plot for splice junction visualization was generated by IGV.

### PCR, reverse transcription (RT)-PCR and RT-qPCR

To validate DNA break sites, total genomic DNA extracted by QIAamp DNA blood mini kit (Qiagen, Gaithersburg, MD) was used in PCR with the following primers: oLLY492/oLLY485 and oLLY489/oLLY495 for CaSki cell DNA, oLLY407/oLLY397 and oLLY402 /oLLY409 for SiHa cell DNA, oLLY469/oXHW48 and oLLY474/oLLY490 for HeLa cell DNA (Table S1). For RT-PCR, total RNA was treated with Turbo DNase (Thermo Fisher Scientific), converted to cDNA by RT with random hexamers and Moloney murine leukemia virus (MuLV) reverse transcriptase (Thermo Fisher Scientific) and used in PCR with the following primers: oZMZ215/oLLY399 and oLLY405/oLLY399 for SiHa cell cDNA and oLLY473/oLLY480 for HeLa cell cDNA (Table S1).

RT-qPCR was carried out with total RNA isolated from SiHa cells using a TaqMan Gene Expression kit (Thermo Fisher Scientific, #4369016). Custom-designed primers and TaqMan probe to specifically detect the viral-host fusion transcripts are listed in Table S1. GAPDH TaqMan probe set (Thermo Fisher Scientific, assay #Hs02786624_g1) was used as an internal control.

### 5’ and 3’ rapid amplification of cDNA ends (RACE)

The 5’ and 3’ RACE assays were conducted with a Smart RACE cDNA amplification kit (CloneTech, Mountain View, CA) as recommended with 1 μg/reaction of total RNA as a template. The following primers (see Table S1) were used: HPV16-specific primer oZMZ260 was used for CaSki cell RNA and HPV18-specific primers oLLY478, oMA119, and oLLY475 were used for HeLa cell RNA.

### Northern blotting

Northern blotting was performed as previously described^45^. Briefly, each 10 μg of total RNA or poly(A)+ mRNA selected from 50 or 100 μg of total RNA was separated in formaldehyde-containing 1% agarose gel in 1× MOPS (morpholinepropanesulfonic acid). The separated RNAs were then capillary transferred onto a nylon membrane and cross-linked by UV. After pre-hybridization, the membranes were probed with ^32^P-labeled oligos overnight at 42°C. The following probes (Table S1) were used for the hybridization: oZMZ302 for detection of HPV16 in CaSki cells; a mixture of oMA196, oZMZ302, and oZMZ220 for detection of HPV16 in SiHa cells; oXHW86 for HPV18 in HeLa cells and oZMZ270 for human GAPDH in all cell lines. The specific signal was captured by Typhoon biomolecular imager (GE Healthcare).

### Cell proliferation and cell cycle distribution assays

The cell proliferation assay based on dehydrogenase activities was performed by Cell Counting Kit-8 (Dojindo Molecular Technologies, Rockville, MD) as recommended. The relative cell viability was calculated based on measured 450-nm absorbance in six biological repeats. For the cell cycle distribution analysis, cells at 1.5 million were fixed with 70% ethanol at 4°C overnight, washed with PBS, followed by the incubation with PBS containing 200 ug/ml RNase A (Thermo Fisher Scientific), 20 ug/ml propidium iodide (Thermo Fisher Scientific), and 0.1% Triton X-100 (Promega) at room temperature for 30 min. Cell cycle distribution was determined by flow cytometry and analyzed by FlowJo software.

### Western blot and antibodies

Total proteins were prepared by direct lysis of 2 million of SiHa cells in 350 ul of 4× SDS-sample buffer and 15 ul of the sample was separated in a NuPAGE Bis-Tris 4– 12% gel (Thermo Fisher Scientific) in 1× MOPS SDS buffer (Thermo Fisher Scientific). After transferred onto a nitrocellulose membrane, the membrane was blocked for 1 h with 5% skimmed milk in 1× TBS (Tris-buffered saline) containing 0.05% Tween (TTBS) and incubated with a primary antibody diluted in TTBS overnight at 4°C. After three washes with 1X TTBS buffer, the membrane was then incubated by a corresponding secondary peroxidase-conjugated antibody diluted in 2% milk/TTBS for 1 h at room temperature. The specific signal on the membrane was generated with SuperSignal West Pico (Thermo Fisher Scientific) and captured by the ChemiDoc Touch imaging system (Bio-Rad).

Anti-p53 (Proteintech, # 10442-1-AP) and anti-beta tubulin (Sigma-Aldrich, #T5201) were used for Western blot.

## Acknowledgements

This work was supported by the Intramural Research Program of the National Institutes of Health, the National Cancer Institute, and the Center for Cancer Research (ZIASC010357 to Z.M.Z.).

## Author contribution

L.Y. and Z.M.Z designed the study, performed all data analyses and interpretations. L.Y. performed all the validation experiments. A.L. and M.C. performed the RNA-seq data analysis. V.M. assisted with data analysis and interpretations. S.M. and V.B. performed SiHa cell RNA sequencing. H.L. performed CaSki cell RNA sequencing. L.Y. and Z.M.Z. wrote the manuscript with input from all co-authors.

## Supplemental legends

**FIG S1.**
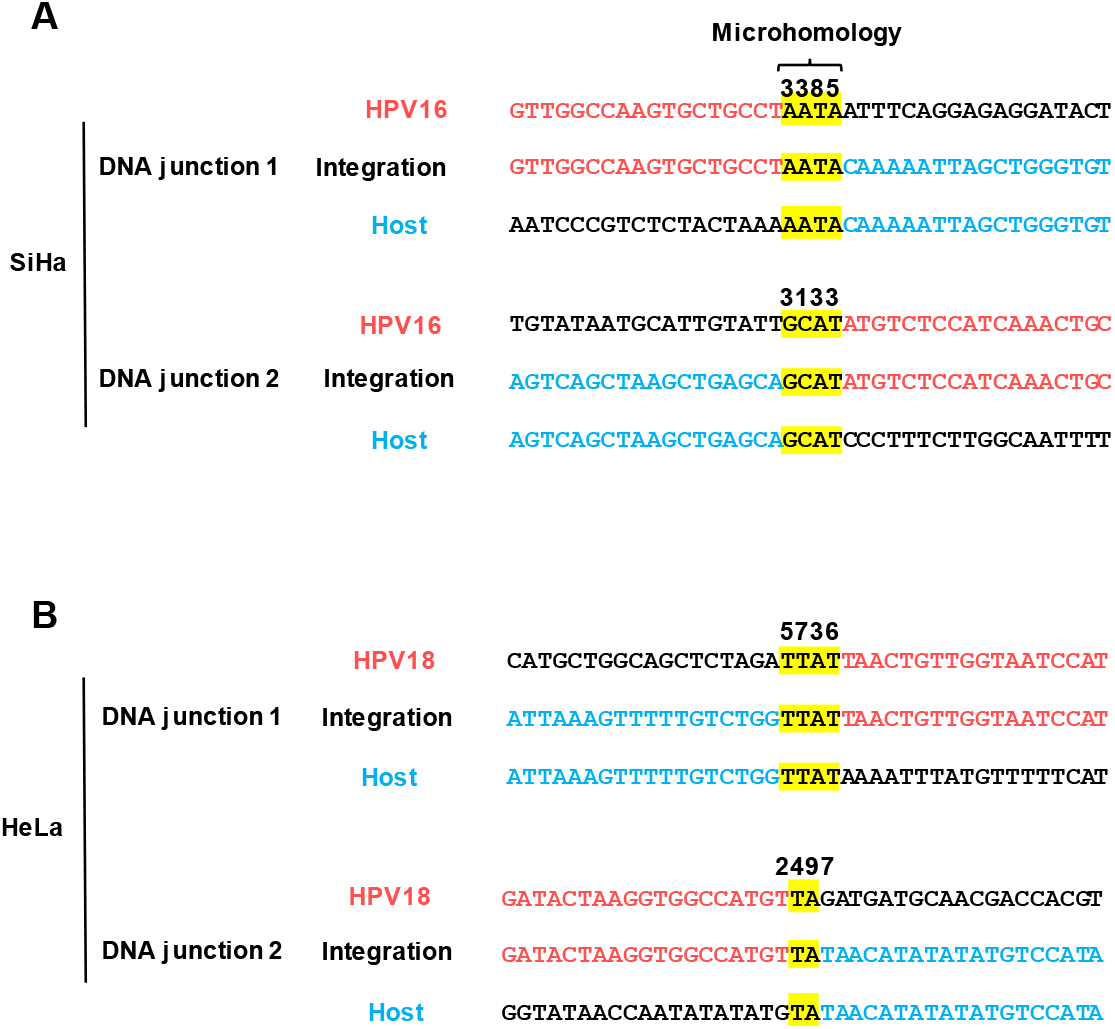
Microhomologous sequence (MHs) motifs identified at virus-host integration junctions in SiHa (**A**) and HeLa (**B**) cells. Yellow color shows the MHs at the integration junction between HPV16/18 and human (hg38) reference genome.

**FIG S2.**
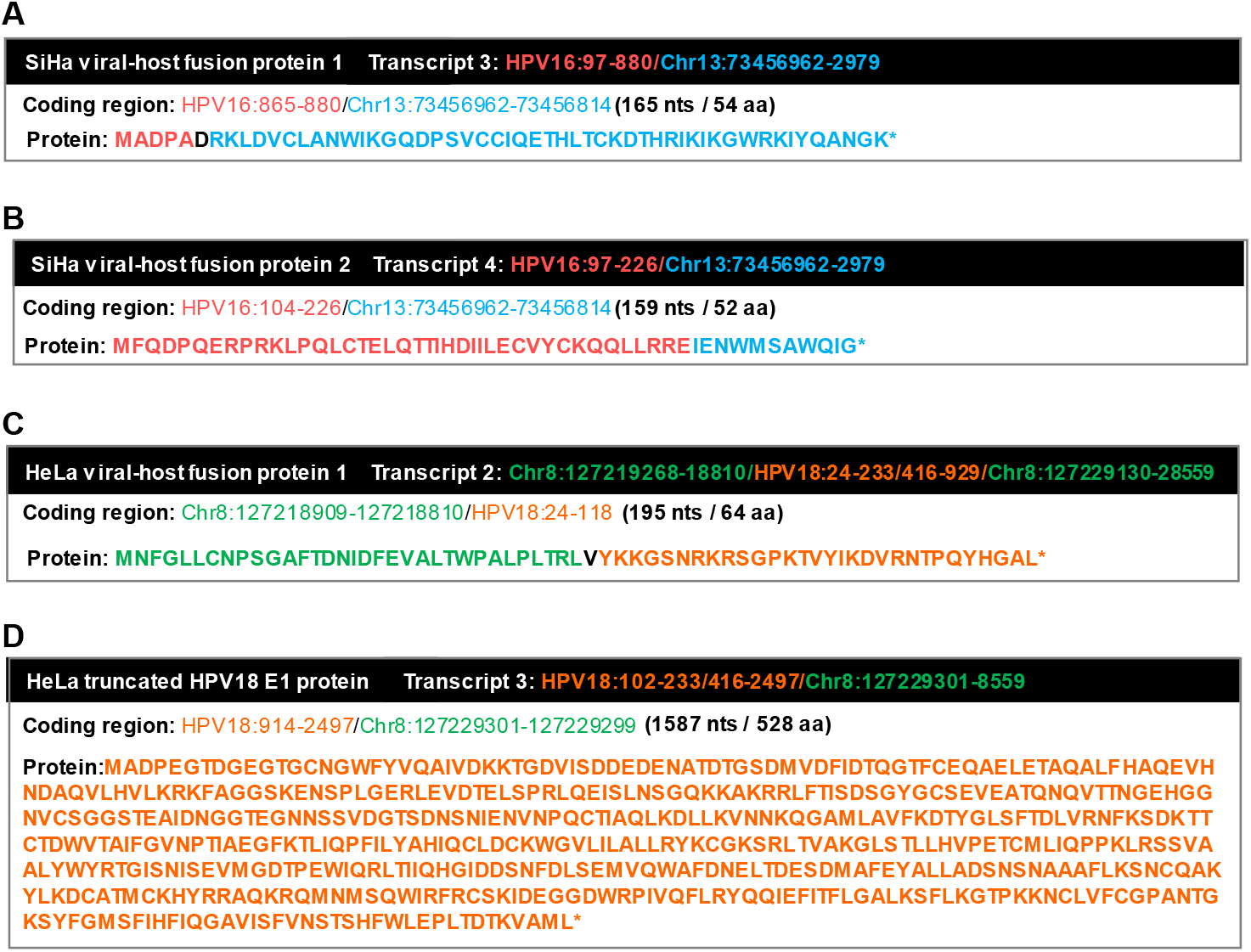
Amino acid sequence of predicted viral-host fusion proteins and truncated E1 protein expressed from chimeric virus-host RNA transcripts detected in SiHa (**A** & **B**) and HeLa (**C** & **D**) cells. The transcript 3 retaining an E1 intron in HeLa cells may encode a truncated E1 protein in size of 528 aa residues by using a host stop codon right from the integration junction. A full size of HPV18 E1 has 658 amino acid residues.

**FIG S3.**
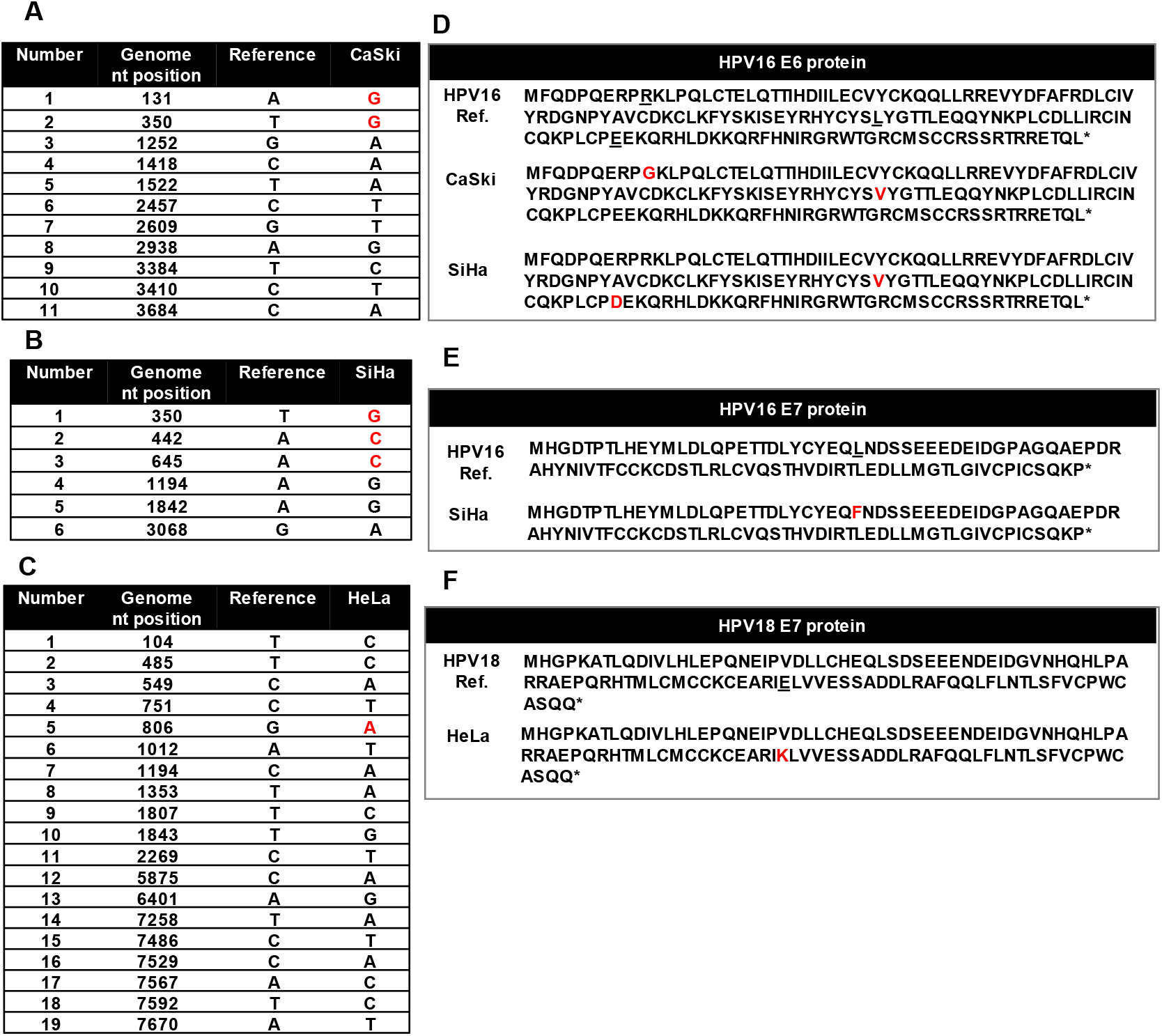
Point mutations in the integrated viral genome in CaSki (**A**), SiHa (**B**) and HeLa (**C**) cells identified by RNA-seq lead to missense mutations in E6 and E7 proteins. Most mutations are silent mutations and only the missense mutations (red color) in the integrated virus genome encode different amino acid residues in E6 and/or E7 proteins in CaSki (**D**), SiHa (**E**) or HeLa (**F**) cells.

## Supplemental Tables

**Table S1.**
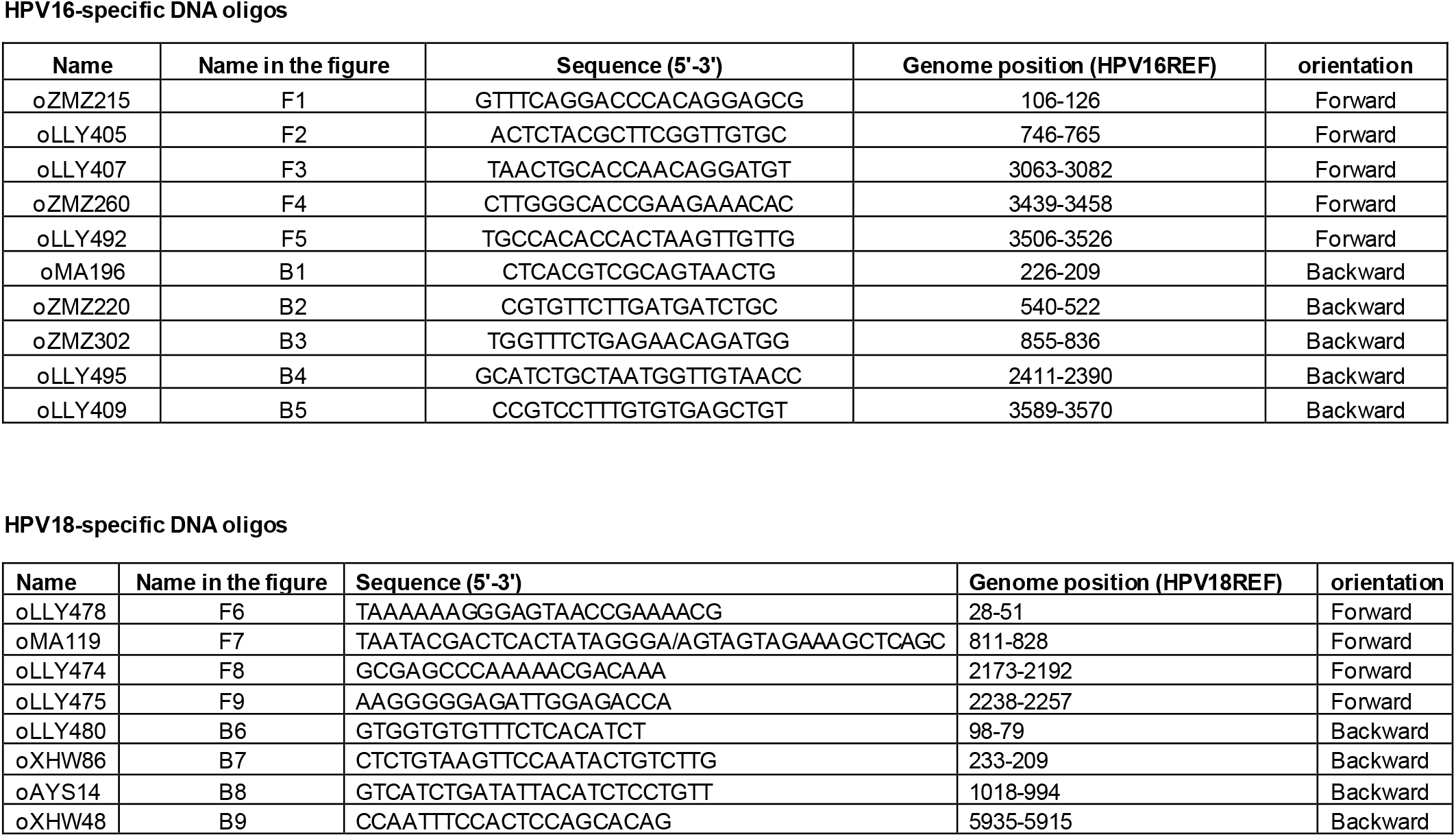
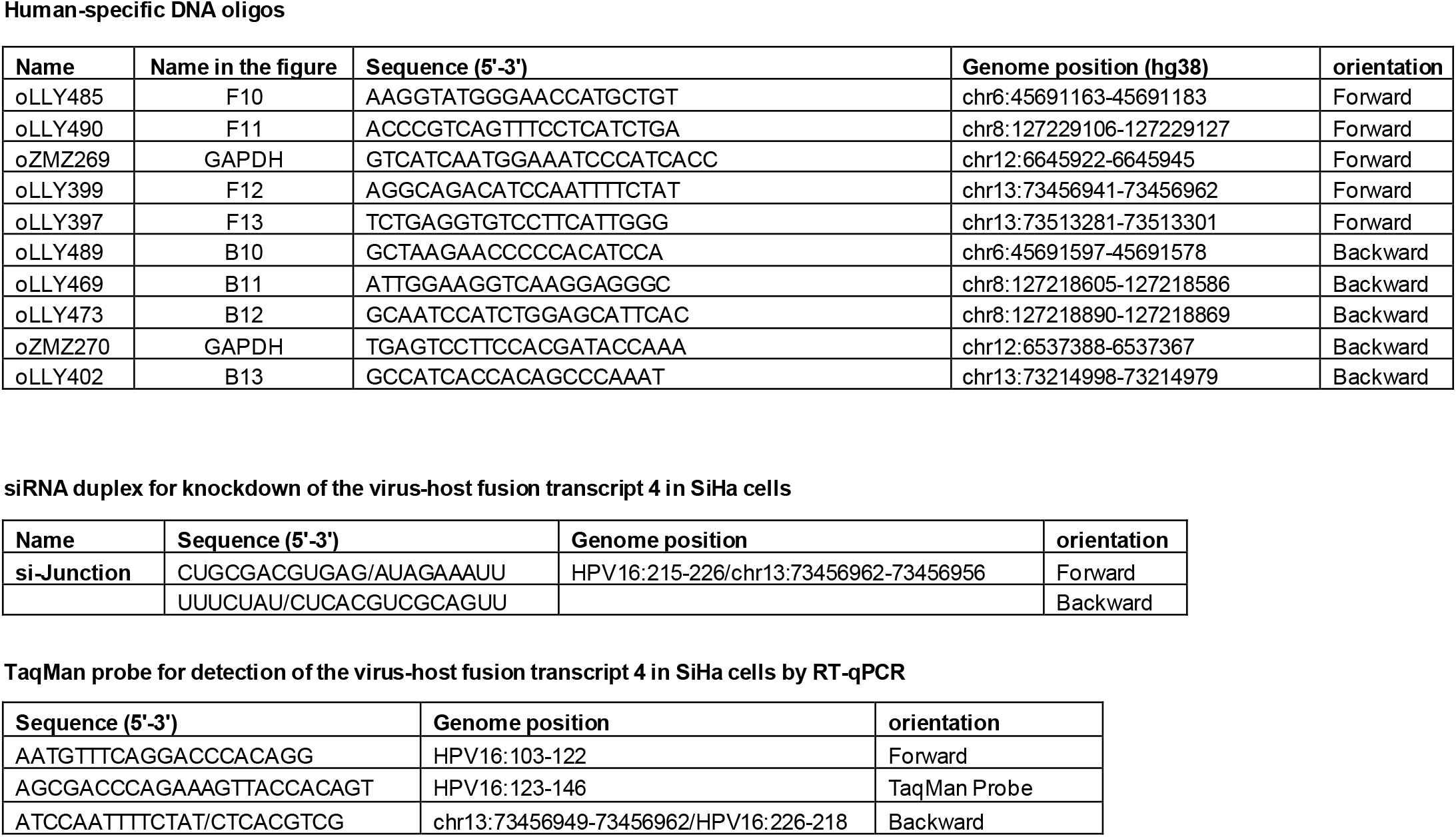
DNA oligos, siRNAs and TaqMan probes used in the study.

## Notes

### Competing Interest Statement

The authors have declared no competing interest.

